# Human antimicrobial peptide, LL-37, induces non-inheritable reduced susceptibility to vancomycin in *Staphylococcus aureus*

**DOI:** 10.1101/2019.12.24.886184

**Authors:** Cathrine Friberg, Jakob Haaber, Martin Vestergaard, Anaëlle Fait, Veronique Perrot, Bruce Levin, Hanne Ingmer

## Abstract

Antimicrobial peptides (AMPs) are central components of the innate immune system providing protection against pathogens. Yet, serum and tissue concentrations vary between individuals and disease conditions. We demonstrate that the human AMP LL-37 lowers the susceptibility to vancomycin in the community-associated methicillin-resistant S. aureus (CA-MRSA) strain FPR3757 (USA300). Vancomycin is used to treat serious MRSA infections, but treatment failures occur despite MRSA strains being tested susceptible according to standard susceptibility methods. Exposure to physiologically relevant concentrations of LL-37 increased the minimum inhibitory concentration (MIC) of S. aureus towards vancomycin by 75% and resulted in shortened lag-phase and increased colony formation at sub-inhibitory concentrations of vancomycin. Computer simulations using a mathematical antibiotic treatment model indicated that a small increase in MIC might decrease the efficacy of vancomycin in clearing an S. aureus infection. This prediction was supported in a Galleria mellonella infection model, where exposure of S. aureus to LL-37 abolished the antimicrobial effect of vancomycin. Thus, physiologically relevant concentrations of LL-37 reduce susceptibility to vancomycin, indicating that tissue and host-specific variations in LL-37 concentrations may influence vancomycin susceptibility in vivo.

## Introduction

Bacterial resistance to antibiotics is generally associated with genetic changes, such as point mutations in the core genome or acquisition of resistance genes present on mobile genetic elements^1^, or intrinsic resistance to antimicrobials, such as impermeability of the outer membrane of Gram-negative bacteria or the active efflux of antimicrobials provided by chromosomally encoded efflux pumps^2^. Previously, we showed that exposure to the synthetic antimicrobial peptide (AMP) colistin reduces susceptibility to vancomycin in the clinically important, methicillin resistant *Staphylococcus aureus* (MRSA) strain USA300 via induction of the GraRS cell wall regulon^3^. *S. aureus* is a human pathogen that causes a variety of diseases and particularly the MRSA clones are becoming widely distributed both in the hospital environment and in the community^4^. MRSA infections are often treated with the glycopeptide antibiotic vancomycin. The CLSI susceptibility breakpoint for vancomycin is 2 μg/ml^5^. However, in the last decade vancomycin-intermediate *S. aureus* strains (VISA) have emerged with reduced vancomycin susceptibility (MIC = 4-8 μg/ml)^6^. These strains are characterized by chromosomal mutations that reduce negative cell wall charge^7^, increase cell wall thickness^8^ or reduce Triton-X mediated autolysis^9^. Even for non-VISA strains vancomycin treatment failures have been linked to strains displaying MIC in the higher end of the susceptibility range (≥1.5 μg/ml)^5,10–13^. Our previous work revealed that upon colistin exposure, *S. aureus* cells transiently and reversibly induce cellular changes that result in a phenotypic VISA-like appearance^3^. This reduced vancomycin susceptibility state will remain undetectable by routine clinical testing and thus the MIC of vancomycin upon colistin exposure *in vivo* may be higher than expected from such testing.

Colistin is a cationic AMP used in treatment of Gram-negative infections, however AMPs are also major components of the innate immune system and a critical defence against a large number of pathogens^14^. Human AMPs are small, often cationic peptides with broad-spectrum antimicrobial activity caused by membrane-damaging pore-formation^14^. An example is the cathelicidin, LL-37^15^. It is expressed by a variety of immune cells, including polymorphonuclear leukocytes (PMN) as well as by epithelial cells and has been identified in saliva, sweat glands, and other tissues^16^. In keratinocytes, exposure to *S. aureus* leads to upregulation of LL-37 production^17^. LL-37 has broad antimicrobial activity against bacteria, fungi and viruses^16^ and in addition is chemotactic for human neutrophils, monocytes and T cells^18^. Plasma levels of LL-37 in healthy individuals have been measured to be approximately 1 μg/ml, but with variations between individuals from 0.5 to 3 μg/ml^19^. In areas of bacterial infection the local concentration will reach even higher levels due to de-granulation of PMNs, e.g. in bronchoalveolar lavage fluid, where LL-37 concentrations from 5 to 25 μg/ml have been measured in infected individuals^20,21^. In the present study, we have examined the interplay between LL-37 and vancomycin and we demonstrate that LL-37 transiently reduces susceptibility to vancomycin. We speculate that natural variation in LL-37 concentrations may lead to variation in susceptibility to vancomycin therapy between tissues and individuals.

## Materials and methods

### Bacterial strains and growth conditions

*S. aureus* USA300 FPR3757^22^ was obtained from the American Type Culture Collection (ATCC). *S. aureus* strains were routinely grown in Mueller-Hinton (MH) or cation adjusted MH (MHII) medium (Sigma) at 37°C in Erlenmeyer flasks with aeration.

### Minimum inhibitory concentration

*S. aureus* FPR3757 was diluted to 10^5^ CFU/ml in MH or MHII media and incubated for 1 h (200 rpm, 37 °C) with either no, 5, 10 or 20 μg/ml LL-37 (Sigma) followed by addition of vancomycin (Sigma). The minimum inhibitory concentration (MIC) was determined in a 96-well microtiter plates (200 rpm, 37 °C, 24 h using technical duplicates and biological triplicates) by visual inspection. Similar approach was used for determining MIC for LL-37 with increasing LL-37 concentrations and no addition of vancomycin.

### Vancomycin susceptibility testing by plating

Exponential phase *S. aureus* FPR3757 cultures were adjusted to OD_600_ = 0.25 in pre-heated MH broth and was then treated with sub-inhibitory concentrations of LL-37 (0, 5, 10, 20 and 40 μg/ml) for 1 h. The cultures were adjusted to OD_600_ = 0.1 before spotting 10 μl aliquots of serial four-fold dilutions on freshly prepared MH agar plates containing increasing concentrations of vancomycin (0, 1, 1.25 and 1.5 μg/ml).

In one experiment, cultures of *S. aureus* FPR3757 were grown with 10 μg/ml LL-37 for 60 min and LL-37 was removed from the medium by spinning (5,000g, 5 min, room temperature) and re-suspended in pre-heated MH broth in a subset of the culture, while exposure to LL-37 was maintained in the other subset. After 0, 60 and 90 min, the cultures where plated on agar plates containing vancomycin (1.5 μg/ml).

The agar plates were inspected for growth following 24 h incubation.

### Growth experiments

Overnight cultures were diluted to approx. 10^5^ CFU/ml in fresh MH broth, and incubated for 1 h at 37°C and 200 rpm. The culture was then split in appropriate aliquots and treated with LL-37 or left untreated for 1 h at 37°C and 200 rpm. The untreated and pre-treated cultures were then split in appropriate aliquots and treated with vancomycin (Sigma) or left untreated. These were then further aliquoted into the wells of a 100-well honeycomb plate and incubated in a Bioscreen C Analyzer (Oy Growth Curves Ab Ltd, Helsinki, Finland) at 37°C and medium continuous shaking.

### Determination of Triton X-100 induced autolysis

The setup of the autolysis assay was conducted as previously described^3^ containing the following changes. Briefly, strains were grown in MH broth at 37°C, 180 rpm, to mid-exponential phase. The cultures were diluted to an OD_600_ of 0.15 in warm MH, grown to an OD_600_ of 0.25, and 10 μg/ml LL-37 was added to one subset of cultures while another subset of cultures served as the unexposed control. After 1 h, the cultured cells were washed twice in ice-cold sterile distilled water and resuspended in the same volume of 0.05 M Tris-HCl (pH 7.2) containing 0.05% Triton X-100. Cells were incubated at 30°C, and the OD_600_ was measured every 30 min. Data are expressed as the percent loss of OD_600_ at the indicated times compared to findings at time zero. Each data point represents the mean and standard deviation from three independent experiments.

### A model of antibiotic treatment

To illustrate the relationship between the MIC of vancomycin and the course of treatment, we used a simple mathematical model of the population dynamics of antibiotic treatment. In this model, we assume Hill function to specify the relationship between the concentration of the antibiotic, A (μg/ml), and the rates per cell per hour at which the exposed bacteria grow and/or are killed ^23,24^.

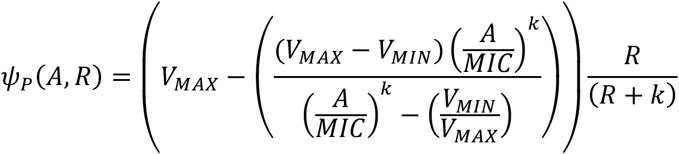

The parameter V_MAX_ > 0 per cell per hour, is the maximum rate of growth of the bacteria in the absence of the antibiotic and V_MIN_ <0 per cell per hour is the minimum rate of growth of those bacteria in the presence of the antibiotic. The parameter κ controls the shape of this function, the greater the value of κ the more acute the relationship between the concentration of the antibiotic and the rate of decline in the viable density of the bacteria^23^. To account for the resource limitation and the fact that as the availability of resources decline the rates at which bacteria grow declines^25^ we assume a hyperbolic function for the relationship between the resource concentration, R (μg/ml), and the rate of growth of the bacteria, R/(R+k), where k, the Monod constant, is the concentration of the limiting resource where, in the absence of the antibiotic the rate of the bacteria is half its maximum value. Thus, when the concentration of the antibiotic and resource are respectively A and R, the growth/death of the bacteria is ψ_P_(A,R) per cell per hour.

We assume the bacteria are maintained in a continuous (chemostat-like) culture of unit volume into which the limiting resources, from a reservoir where it is maintained at a concentration C (μg/ml), flow in at a constant rate w (ml per hour) with the same rate at which excess resource, bacteria and antibiotics flow out^26^. The bacteria take up the limiting resource at rates proportional to their maximum growth rates and a conversion efficiency parameter, e (μg/cell). With these definitions and assumptions, the rates of change in the density of viable bacteria, P (cells per ml), the limiting resource and antibiotics are expressed as a set of coupled differential equations.

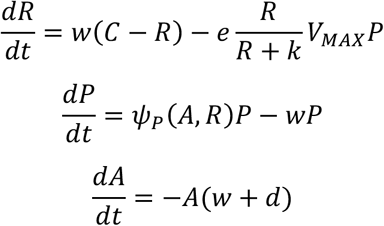

where d is the rate of decay in the concentration of the antibiotic.

To simulate the dynamics of treatment, we solve these equations numerically using Berkeley Madonna. Copies of this program are available from. To account for treatment, in these simulations, an antibiotic at a concentration of A_MAX_ (μg/ml) is introduced into the vessel every *Dose* hours.

### Galleria mellonella infection model

Twenty healthy 5th instar larvae each weighing 250 mg were injected with 2.5 x 10^6^ CFU of *S. aureus* FPR3757 followed by 10 μg/ml LL-37 or 0.9 % NaCl for controls. One hour post infection, the larvae were treated with 15 mg/kg bodyweight of vancomycin, corresponding to a relevant clinical dose. The LL-37 pre-treatment and antimicrobial therapy was repeated every 24 h for a total of 72 h and survival was monitored. Controls for toxicity of LL-37 combined with the antimicrobials as well as for non-treated *S. aureus* infection were included with all larvae receiving an equal number of injections of either active compound or 0.9 % NaCl.

### Statistics

The data were analyzed in GraphPad Prism 7 (GraphPad Software Inc.) using one-way analysis of variance (ANOVA) with a *post hoc* analysis of Dunnett’s multiple comparison tests, where *P < 0.05* (*), *P < 0.01* (**) and *P < 0.001* (***). The survival data was analyzed by using the Kaplan-Meier method and statistical difference determined, using log-rank test.

## Results

### Pre-exposure to LL-37 reduces S. aureus susceptibility to vancomycin

We have previously shown that pre-exposure of *S. aureus* to the antimicrobial peptide, colistin, reduces vancomycin susceptibility^3^. To address if exposure to the human antimicrobial peptide LL-37 also affects vancomycin susceptibility we initially determined the LL-37 MIC of *S. aureus* FPR3757 to be >128 μg/ml. As concentrations of LL-37 in the range of 0-40 μg/ml are physiologically relevant we pre-treated cells with either 10 or 20 μg/ml LL-37 prior to addition of vancomycin. By this approach, we found that in MH the vancomycin MIC increased from 0.8 μg/ml to 1.2 and 1.4 μg/ml, respectively, upon exposure to either 10 or 20 μg/ml LL-37. In MHII the increase was from 1.2 μg/ml to 1.5 μg/ml following pre-treatment with 20 μg/ml LL-37. We also examined the effect of 5 μg/ml LL-37 and found that it did not impact vancomycin susceptibility. To further examine the effect of LL-37 on vancomycin susceptibility, we spotted serial dilutions of cultures pre-treated with various sub-inhibitory concentrations of LL-37 on agar plates containing increasing concentrations of vancomycin. Cultures exposed to 40 μg/ml LL-37 were able to form colonies in the presence of 1.5 μg/ml vancomycin in contrast to untreated cells, where no growth was observed (Figure 1).

**Figure 1.**
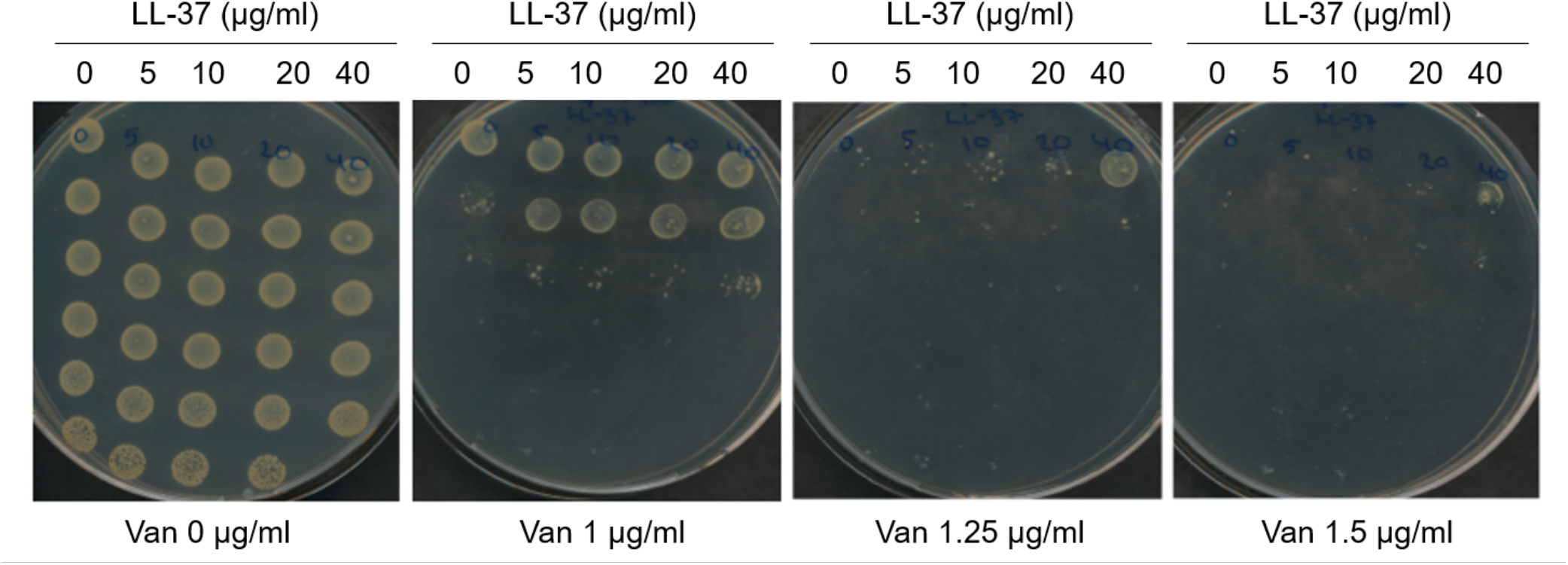
LL-37 influences vancomycin susceptibility of *S. aureus*. Exponential phase *S. aureus* FPR3757 cultures were pretreated with 0, 5, 10, 20 and 40 μg/ml LL-37 for 1 h before spotting serial four-fold dilutions on MH agar containing increasing concentrations of vancomycin.

When observing the growth dynamics of treated and untreated cells in liquid cultures, pre-treatment with LL-37 notably reduced the vancomycin induced lag-phase measured as the time from inoculation until density reached OD_600_ = 0.1 (Figure 2a and b). In the presence of vancomycin (at 0.75 μg/ml) the lag-phase was 24.5 (±1.9) h for untreated cells, whereas the lag-phase was 8.1 (±2.5) h following pre-exposure to 20 μg/ml LL-37 (p<0.001). Shortening of the lag phase was proportional to the concentration of LL-37 used for pre-treatment (Figure 2a and b). In the absence of vancomycin, LL-37 treatment did not affect growth kinetics (Figure 2a). When determining maximum growth rate, no effect of LL-37 was observed in the vancomycin-treated or control cultures (Figure S1), indicating that shortening of lag-phase rather than growth rate is important for the LL-37 induced reduction in vancomycin susceptibility.

**Figure 2.**
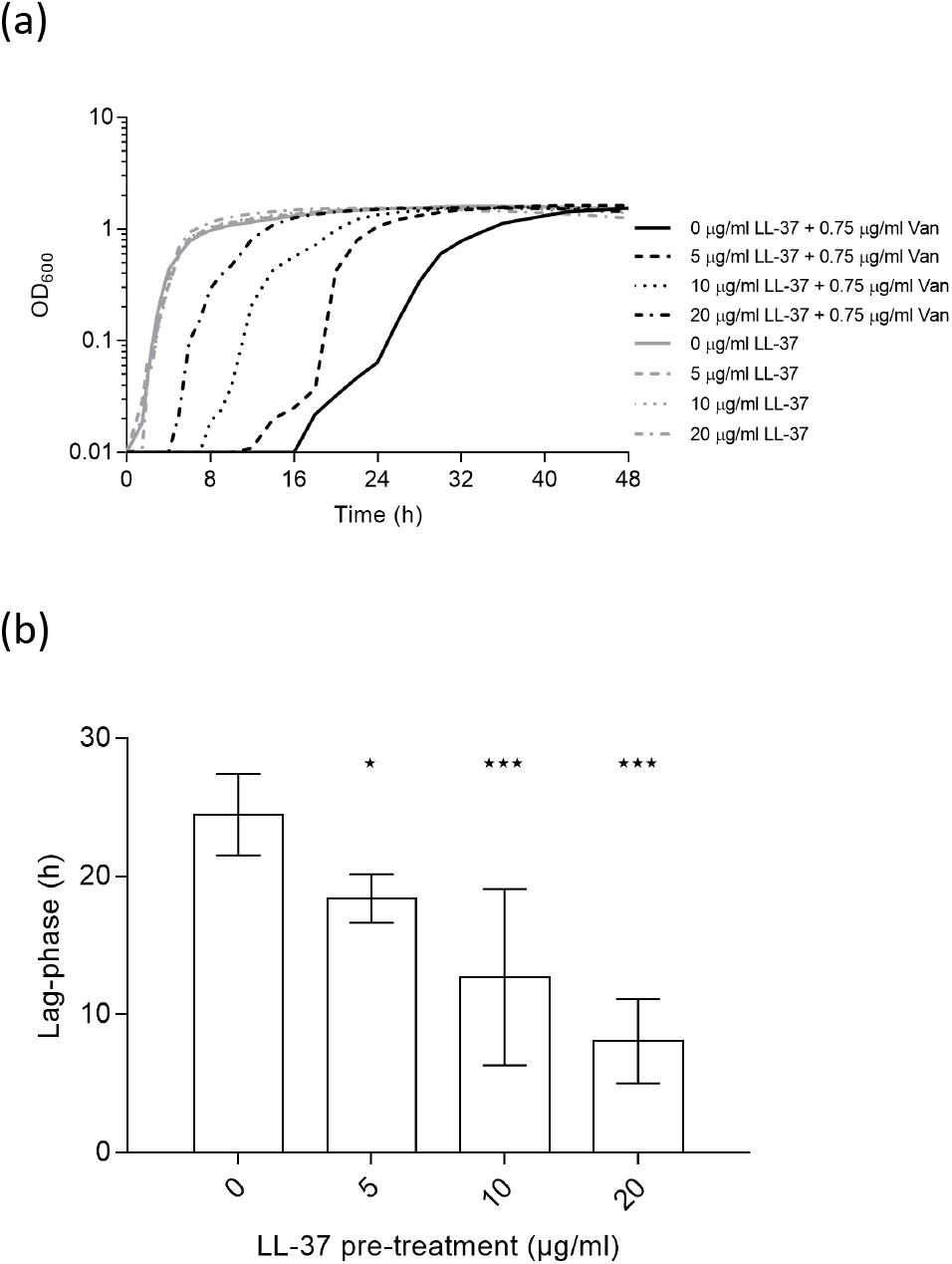
LL-37 influences vancomycin susceptibility of *S. aureus* in liquid culture. (a) The growth of S. aureus FPR3757 with (black lines) or without (grey lines) exposure to 0.75 μg/ml vancomycin after treatment for 1 h with 0, 5, 10 or 20 μg/ml LL-37. The curves are an average of five replicates. (b) The lag-phase of the same cultures depicted in (a) (n=5). Error bars display 95% confidence intervals. **P<0.01; *P<0.05

### LL-37 induces reversible vancomycin tolerance

To examine if continuous LL-37 exposure is needed for the reduced vancomycin susceptibility we monitored the LL-37 exposure time required before being able to observe reduced susceptibility to vancomycin and how long the cells retained the reduced susceptibility after removal of LL-37. We found that the phenotype was observed already after 10 minutes of exposure to LL-37 and that it lasted approximately 60 minutes after LL-37 was removed from the growth medium (Figure 3). These results demonstrate that the reduced vancomycin susceptibility induced by LL-37 exposure is transient and non-inherited.

**Figure 3.**
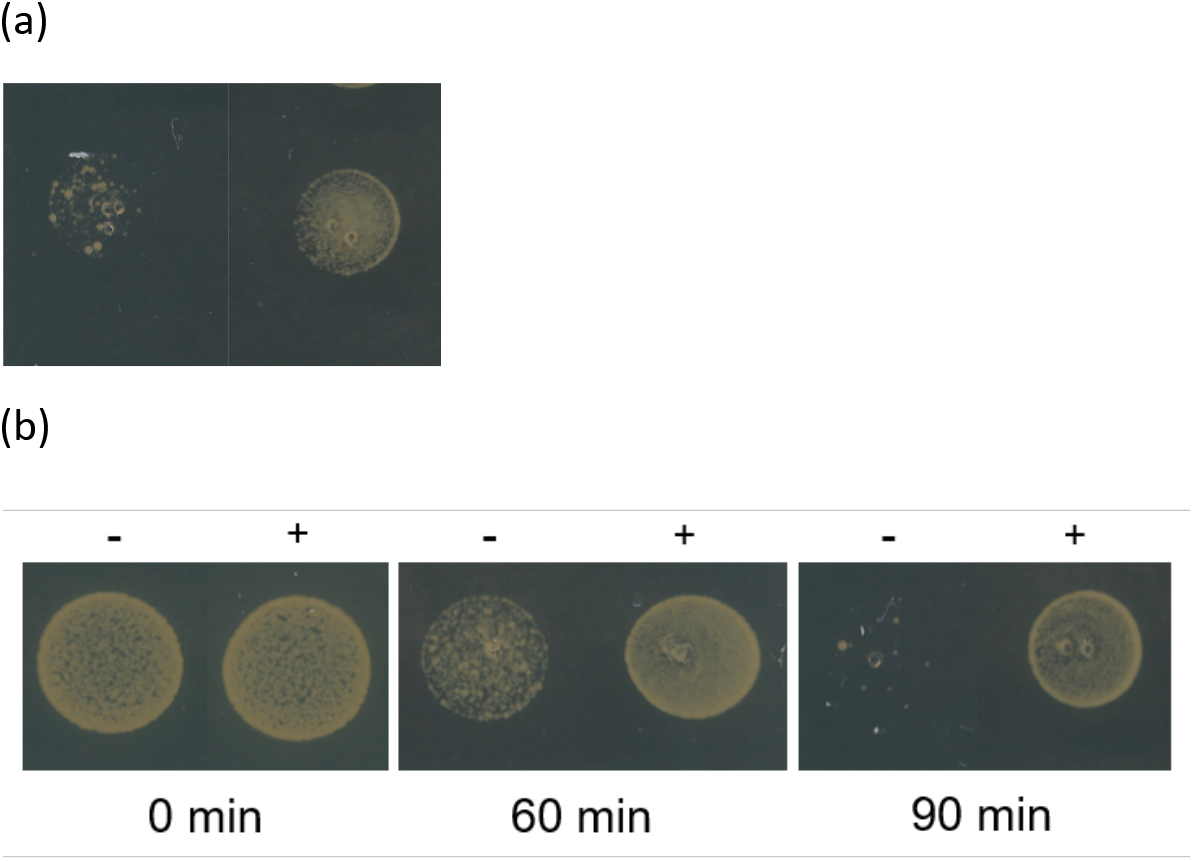
Transient induction of reduced vancomycin susceptibility by LL-37. (a) Cultures of *S. aureus* FPR3757 were grown without (left) or with (right) 10 μg/ml LL-37 for 10 minutes and spotted on vancomycin plates (1.5 μg/ml). (b) Cultures of *S. aureus* FPR3757 were grown with 10 μg/ml LL-37 for 60 min and LL-37 was removed from the medium in a subset of the culture (left), while exposure to LL-37 was maintained in the other subset (right). After 0, 60 and 90 min, the cultures where plated on agar plates containing vancomycin (1.5 μg/ml).

### Effect of LL-37 on autolysis

Reduced susceptibility to vancomycin in genetically stable VISA strains are characterized by an altered cell wall structure that phenotypically can be measured as a decrease in Triton X-100 induced autolysis^9^. We examined the effect of LL-37 exposure (10 μg/ml) on autolysis as described previously^3,7^. Compared to an un-treated control, the pre-treated culture showed a small but noticeable decrease in Triton X-100 induced autolysis (Figure 4) suggesting that LL-37 may impact vancomycin susceptibility via changes in the cell wall.

**Figure 4.**
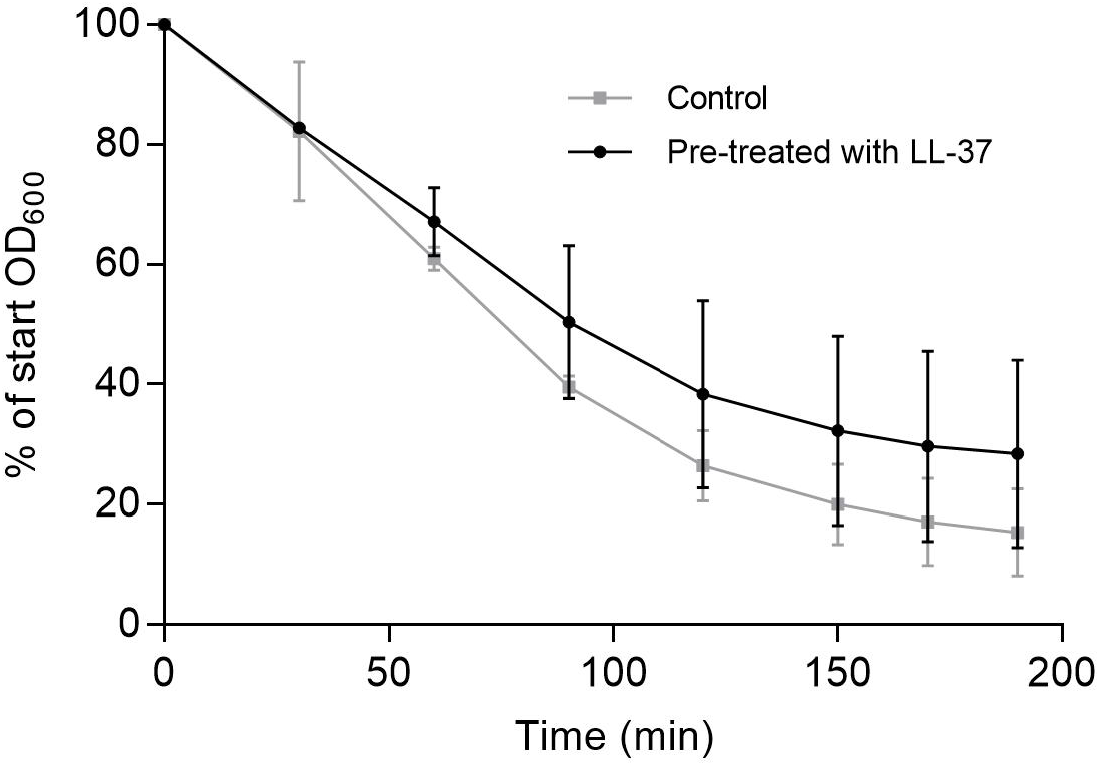
LL-37 exposure reduces Triton X-100 mediated autolysis. LL-37 induced autolysis measured as reduction of bacterial density over time in a 0.05% Triton X-100 solution containing treated (10 μg/ml LL-37) or untreated cultures. Error bars display 95% confidence intervals, (n=3).

### The anticipated relationship between the MIC of vancomycin and the course of treatment

Clinical isolates of vancomycin susceptible MRSA strains with MICs in the higher end of the susceptibility range (≥1.5 μg/ml) are linked to vancomycin treatment failure^5^. Such treatment failures may be related to unidentified host factors, like LL-37, which are causing an increase in *S. aureus* vancomycin MIC^27^. To illustrate how, as suggested by clinical data^5^, small increases in the MIC of vancomycin may lead to treatment failure, we simulated the population dynamics of treatment with the mathematical model described in the Materials and Methods (Figure 5). For any given concentration of the antibiotic, the rate at which the bacteria are killed declines with the MIC of the antibiotic. While treatment with 2.5 μg/ml will clear the bacteria when the MIC is 0.8, it will not when the MIC is 1.4, and will only be manifest as a slow decline in the density of the population when the MIC is 1.4. To be sure, higher concentrations of the antibiotic will be effective in clearing the infection when the MIC is 1.4, but because of toxicity, nephrotoxicity in the case of vancomycin^28^, increasing doses of the drug is not a clinically amenable solution. Stated in another way, *S. aureus* exposed to clinically relevant LL-37 concentrations may have a negative impact on the efficacy of vancomycin for treating infections.

**Figure 5.**
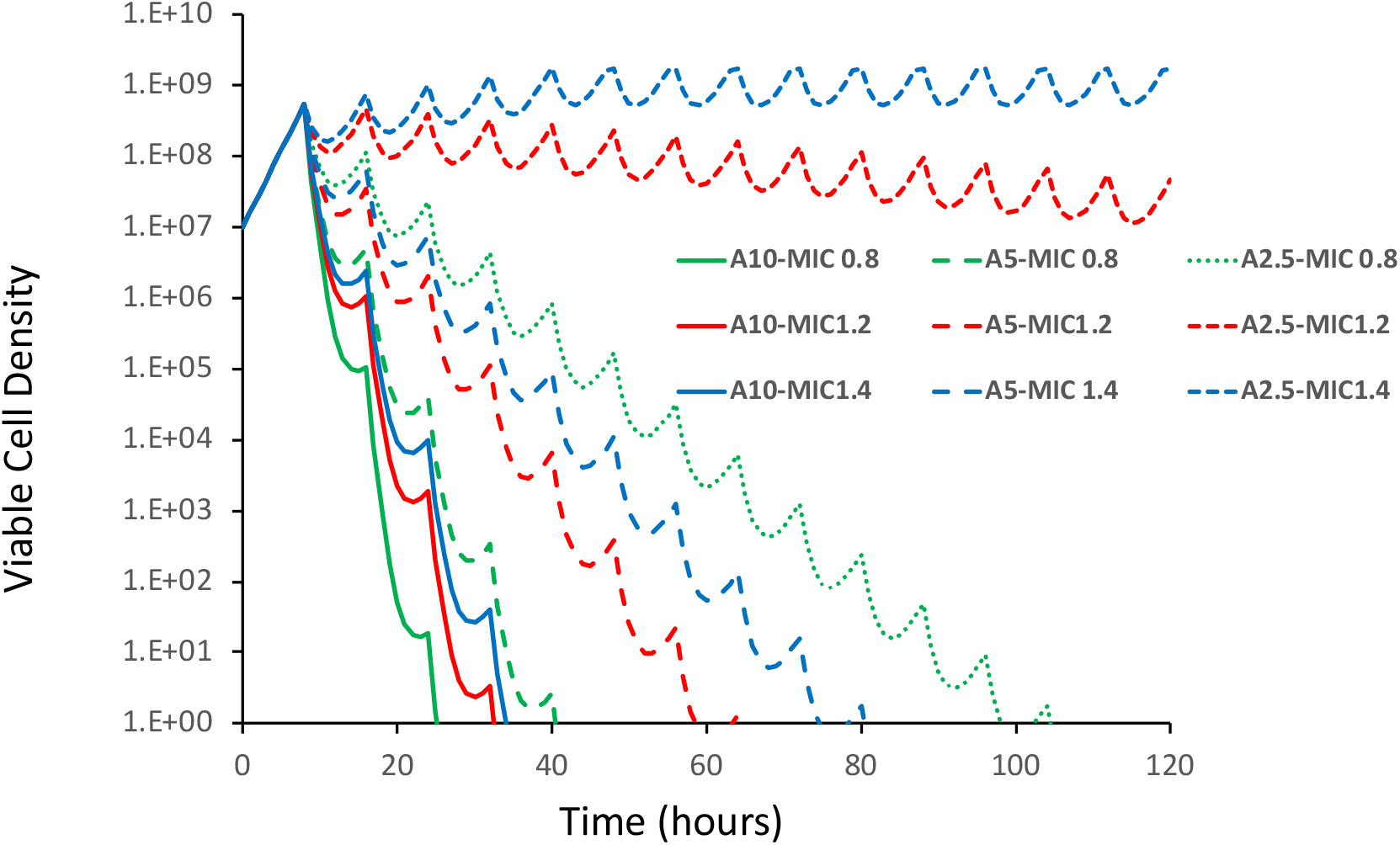
Simulation results of the relationship between the dose of the antibiotic and MIC. Simulations were made of treatment every 8 hours with a drug (10, 5 and 2.5 μg/ml) of cells with the MIC of 0.8, 1.2 and 1.5 μg/ml of the drug and the changes in the viable cell density observed. Hill function parameters V_MAX_= 1.0, V_MIN_=-3.0, κ=1.0, continuous culture parameters, k=1, e=5×10^−7^, w=0.5 and C=1000 (see the equations in the Materials and Methods. See supplemental Figure S2 for the pharmacodynamic Hill functions and the rates of kill bacteria exposed to antibiotics with these Hill function parameters.

### LL-37 abolishes antimicrobial effect of vancomycin in Galleria mellonella

The observed *in vitro* reduction in vancomycin susceptibility after LL-37 exposure combined with the predictions of treatment failure associated with modest increments in MIC from our computer model prompted us to test the efficacy of vancomycin in treating *Galleria mellonella* wax moth larvae^29^ infected with *S. aureus* FPR3757 in the presence or absence of LL-37. In the absence of LL-37, vancomycin reduced the mortality inflicted by the MRSA strain. However, in the presence of LL-37, the antimicrobial potential of vancomycin was completely abolished (P<0.001), rendering the larvae as susceptible to *S. aureus* killing as the non-treated *S. aureus* infection (Figure 6).

**Figure 6.**
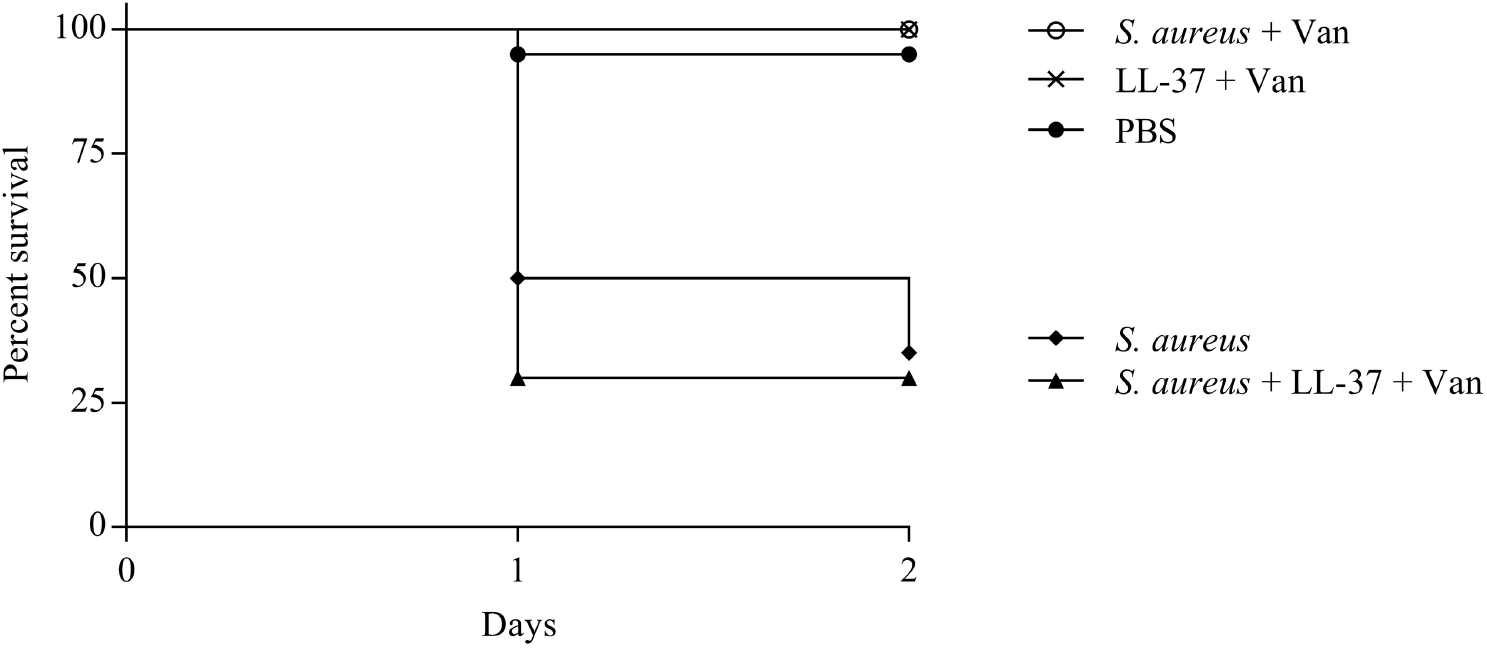
Influence of LL-37 on vancomycin treatment of *S. aureus* infecting *G. mellonella*. Survival of *G. mellonella* wax moth larvae infected with *S. aureus* FPR3757 and with or without LL-37 (10 μg/ml) exposure followed by treatment with vancomycin (15 mg/kg bodyweight). Controls for non-treated *S. aureus* infection and non-infected larvae exposed to combinations of the drugs were included. All larvae received same number of injections. n=20 larvae for each condition.

## Discussion

Antimicrobial chemotherapy is currently based on standardized *in vitro* antimicrobial susceptibility testing, however the assay can face limitations in predicting *in vivo* treatment efficacy^30^. The lack of predictive value is partly attributed to the fact that the growth media in which the susceptibility tests are conducted, do not sufficiently represent the *in vivo* environment. Several factors, such as nutrient availability, oxygen level, pH and presence of innate immune components, have been identified to affect antibiotic susceptibility in both Gram-positive and –negative species^31–34^. Here we show that the presence of the antimicrobial peptide LL-37 of the innate immune response antagonizes the activity of vancomycin against *S. aureus in vitro*. This effect is transient and is also evident in the *G. mellonella* infection model, where the presence of LL-37 with vancomycin significantly reduces survival of *G. mellonella* larvae compared with vancomycin treatment alone. Even though only one strain has been investigated and there may be strain to strain variations, our data suggest that the MIC values reported from *in vitro* assays may underestimate the vancomycin MIC *in vivo* in the presence of LL-37.

Vancomycin is one of the most important antibiotics for treatment of MRSA infections^35^. The CLSI susceptibility breakpoint for vancomycin is 2 μg/ml^5^. Treatment failures reported for infections with MRSA strains being susceptible to vancomycin has led to discussions whether the MIC breakpoint at 2 μg/ml should be further decreased^5^. A meta-analysis associated high-MIC (≥ 1.5 μg/ml) MRSA isolates with increased risk of mortality compared with low-MIC (< 1.5 μg/ml) MRSA isolates^5^, thus indicating that even minor increases in vancomycin MIC can negatively affect treatment outcome and survival of patients with MRSA infections.

To illustrate how small increases in the MIC of vancomycin can affect treatment outcome due to interaction between this drug and LL-37, we used simple mathematical models of the pharmacodynamics of antibiotics and bacteria and the course of antibiotic treatment. While the parameter values chosen for this illustration are in the range estimated for *S. aureus* exposed to vancomycin *in vitro*, these simple simulations certainly do not capture the physical, physiologically, and pharmacological heterogeneity of vancomycin treatment of *S. aureus* in an infected host, much less the contribution of the host’s innate and adaptive immune system to the course of treatment^36,37^. Then again, neither do MICs estimated under *in vitro* conditions, but in combination they may add an additional layer of understanding to the complexity of antimicrobial therapy.

The impact of LL-37 on vancomycin susceptibility under *in vivo* conditions may vary from person to person. As mentioned earlier there is natural variation in the LL-37 levels in healthy individuals that will be increased in sites of infection. Furthermore, LL-37 levels are also influenced by disease conditions, where for example psoriasis patients have increased LL-37 levels^38^, while atopic dermatitis patients have reduced levels^39^. Thus, there is ample evidence for considerable variations in LL-37 levels both from tissue-to-tissue and from person-to-person. Here we provide evidence that such variation may influence susceptibility to vancomycin and our results stress the need for taking host factors into account when testing susceptibility to antimicrobials.

Collectively, our results indicate that a better understanding of the host’s role in reduced vancomycin susceptibility could be important when designing treatment regimens to improve clinical outcome.

## Acknowledgement

This research was funded by grants from the U.S. National Institutes of General Medical Science GM098175-17 and 1R35 GM136407-01 to B.R. L.; from the Danish Council for Independent Research, Technology and Production to J.H. (12-126289) and H.I. (7017-00079B) and the European Union Horizon 2020 research and innovation programme under the Marie Sklodowska-Curie grant agreement No. 765147 to A. F. and H.I.

The funding sources had no influence on study design, data analysis or decision to submit the article for publication.

C.F., J.H., B.L. and H.I. conceived and designed the study. Experiments were performed by C.F., A.F. and J.H. Analysis of data and drafting of the manuscript was performed by C.F., J.H., M.V., V.P., B.L. and H.I. All authors read and approved the final manuscript.

## Ethical approval

Not required.

## Supplementary figures

**Figure S1.**
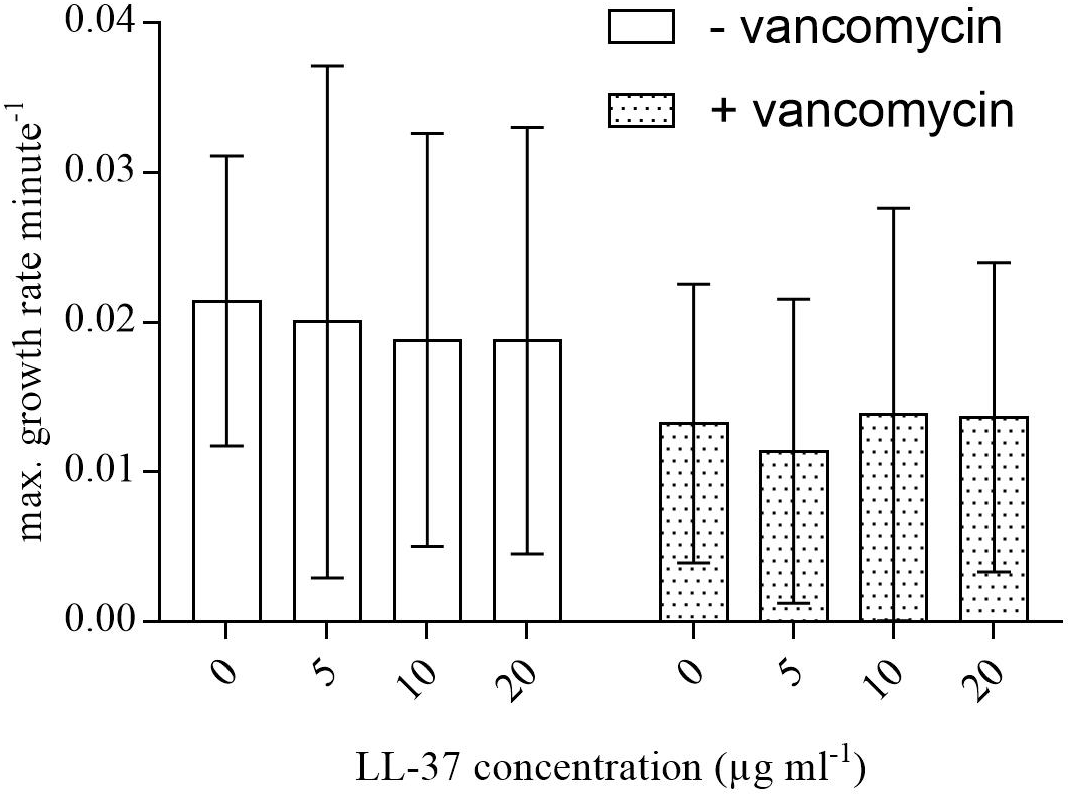
LL-37 effect on maximum growth rate. From the growth experiments as described in Materials and Methods, the maximum growth rates were determined for *S. aureus* cultures pre-exposed to LL-37 (0 to 20 μg/ml) for 1 h before being exposed to 0 or 0.75 μg/ml vancomycin. Error bars represent s.d. of 3 biological replicates each containing 3 technical replicates.

**Figure S2.**
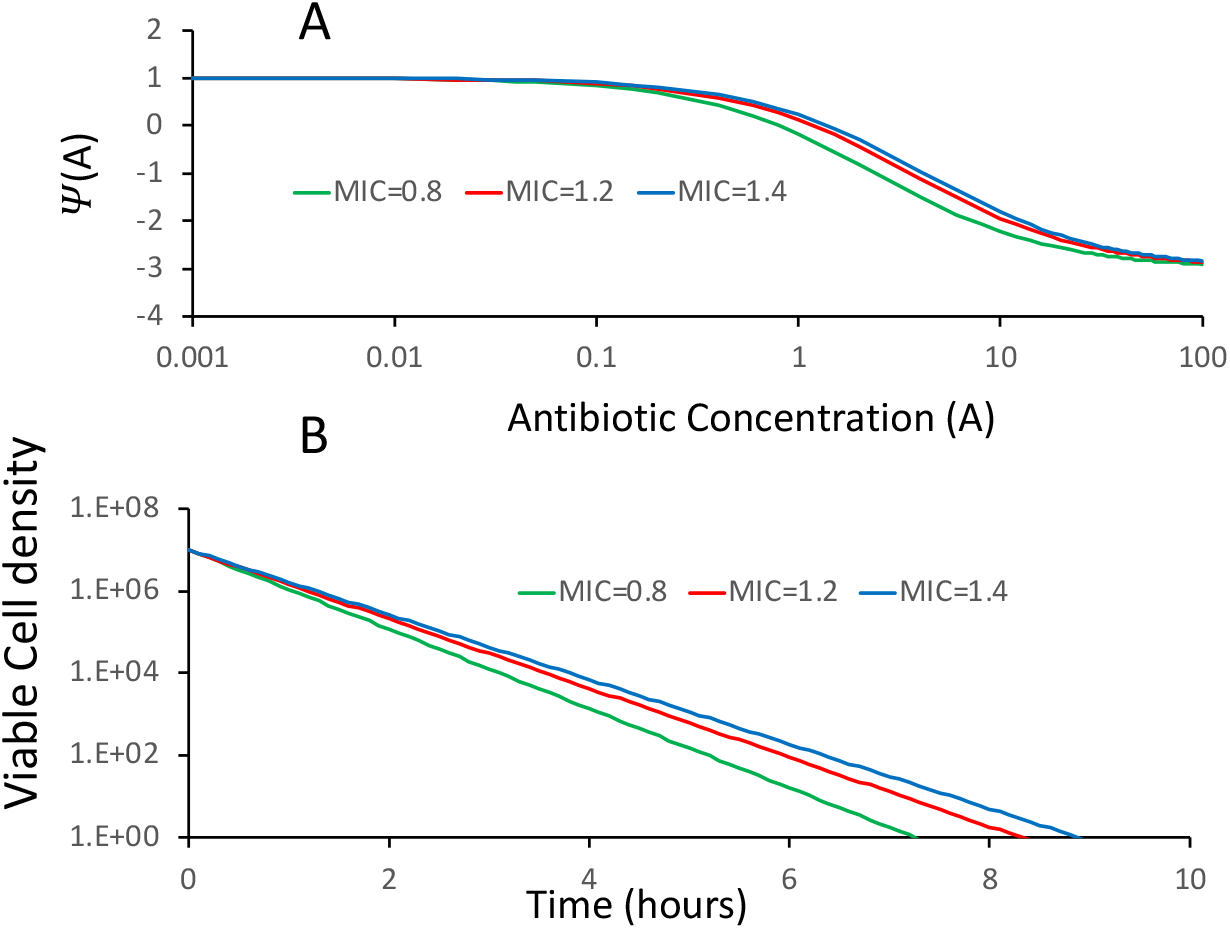
A) Hill function pharmacodynamics and B) the relationship between the concentration of the antibiotic and the rate of kill of bacteria with an antibiotic. The parameters of these functions and kill dynamics are the same as those in used in Figure 5 to illustrate the treatment dynamics anticipated with different concentrations of antibiotics and the MICs of these drugs.

